# High speed large scale automated isolation of SARS-CoV-2 from clinical samples using miniaturized co-culture coupled with high content screening

**DOI:** 10.1101/2020.05.14.097295

**Authors:** Rania Francis, Marion Le Bideau, Priscilla Jardot, Clio Grimaldier, Didier Raoult, Jacques Yaacoub Bou Khalil, Bernard La Scola

## Abstract

SARS-CoV-2, a novel coronavirus infecting humans, is responsible for the current COVID-19 global pandemic. If several strains could be isolated worldwide, especially for *in-vitro* drug susceptibility testing and vaccine development, few laboratories routinely isolate SARS-CoV-2. This is due to the fact that the current co-culture strategy is highly time consuming and requires working in a biosafety level 3 laboratory. In this work, we present a new strategy based on high content screening automated microscopy (HCS) allowing large scale isolation of SARS-CoV-2 from clinical samples in 1 week. A randomized panel of 104 samples, including 72 tested positive by RT-PCR and 32 tested negative, were processed with our HCS procedure and were compared to the classical isolation procedure. Isolation rate was 43 % with both strategies on RT-PCR positive samples, and was correlated with the initial RNA viral load in the samples, where we obtained a positivity threshold of 27 Ct. Co-culture delays were shorter with HCS strategy, where 80 % of the positive samples were recovered by the third day of co-culture, as compared to only 25 % with the classic strategy. Moreover, only the HCS strategy allowed us to recover all the positive elements after 1 week of co-culture. This system allows rapid and automated screening of clinical samples with minimal operator work load, thus reducing the risks of contamination.

## Introduction

An outbreak caused by a novel coronavirus (SARS-CoV-2) has broken in late December 2019 in Wuhan, China, then spread worldwide and was declared a pandemic by WHO on the 12^th^ of March 2020(1),(2),(3). This global health crisis has drawn the attention of the entire scientific community who are working altogether to understand the reason of this outbreak and to find a solution at the levels of rapid diagnosis and effective treatment(4). Several known drugs have been repurposed to treat COVID-19 patients and have shown *in-vitro* and *in-vivo* efficiency(5),(6),(7),(8),(9),(10). Moreover, vaccine development is ongoing in several countries around the world(11),(12), in addition to potential plasma therapy(13),(14). Laboratory diagnosis is mainly based on molecular biology using specific RT-PCR systems to detect the virus in clinical samples(15),(16),(17). However, during such pandemics, strain isolation is important, as having the particle represents the key to all *in-vitro* research such as drug susceptibility testing and vaccine development. Furthermore, culture allows access to all viral genomes since whole genome sequencing techniques performed directly on samples have their limitations in terms of sensitivity.

If routine culture was progressively abandoned in most virology laboratories, we believe that isolating as many strains as possible allows to compare genomic sequences with phenotype of infection, *in vitro* and *in vivo.* This would help understanding the epidemiological aspects of this illness, its physiopathology and better target treatment and prevention(18). A first application of this strategy was used by our group to evaluate the risk of contagiousness of patients for discharge from infectious diseases ward(19). However, the current co-culture strategy is tedious and time consuming, especially due to the large number of samples to be cultured. During the current COVID-19 outbreak, the samples without any observable cytopathogenic effects after 1 week of co-culture, were sub-cultured in blind and monitored for 3 weeks. The best would be to have an automated system allowing the rapid screening and monitoring of co-cultures at large scale. In previous works, we developed a screening strategy based on high content screening microscopy (HCS) for the isolation of environmental giant viruses in amoeba and the strict intracellular bacterium *Coxiella burnetii*(20),(21). In this work, we used the same automated high-throughput method and adapted it for SARS-CoV-2 isolation from clinical samples with the objective to discard the negative co-cultures after 1 week and omit blind sub-cultures. Specific algorithms were applied to detect cytopathic effects in co-cultures at high throughput, which eliminates the subjectivity related to manual observations by the laboratory personnel. This strategy exhibited a similar isolation rate, but a lower co-culture delay when compared to the classic technique routinely used for isolation, as we were able to detect all positive co-cultures in one week.

## Materials and Methods

### 1. Co-culture process for the developmental stage

For protocol development, we used Vero E6 cells (ATCC CRL-1586) as cellular support and the locally isolated SARS-Cov-2 strain IHUMI-3. This viral strain was previously isolated in our lab from a nasopharyngeal swab as previously described(6). The viral titer was calculated by the TCID50 method. Briefly, we cultured Vero E6 cells in black 96-well microplates with optical-bottom (Nunc, Thermo Fischer) at a concentration of 2×10^5^ cells/ml and a volume of 200 µl per well, in a transparent MEM medium supplemented with 4% fetal calf serum and 1% glutamine. Plates were incubated for 24 hours at 37°C in a 5 % CO_2_ atmosphere to allow cell adhesion. Infection was then carried out with 50 µl of the viral stock suspension diluted up to 10^−10^. The plates were centrifuged for 1 hour at 700 x g and the total volume per well was adjusted to 250 µl with culture medium. Uninfected cells were considered negative control.

### 2. Detection process optimization

DNA staining was performed with NucBlue™ Live ReadyProbes™ reagent (Molecular Probes, Life Technologies, USA). A concentration of 4 ng/ml was used (equivalent to 10 µl per well directly from stock solution) and a different well was stained each day to avoid photo-bleaching and possible cytotoxicity, as previously described(21).

Image acquisition and analysis were performed using the automated CellInsight™ CX7 High Content Analysis Platform coupled with an automation system including an Orbitor™ RS Microplate mover and an incubator Cytomat™ 2C-LIN (Thermo Scientific). The HCS Studio 3.1 software was used to set up acquisition parameters using a 20x objective (0.45 NA), and to define image analysis. Autofocus was performed on the fluorescence channel of the fluorescent probe NucBlue (386 nm). This channel served as a primary mask for cell detection and identification. The regions of interest (ROI) were then identified on brightfield images as a Voronoi diagram derived from nuclear masks. Cell debris were removed using area cutoffs. The entire well (80 fields per well) was screened on a daily basis and data were extracted and analyzed in a dedicated application that we recently developed in R Studio® for the detection of the intracellular bacteria, *Coxiella burnetii*(21). We optimized this application for the detection of cytopathic effects caused by Covid-19.

Briefly, a database consisting of negative (uninfected cells) and positive (infected cells) controls was generated. The data were used to define specific features allowing the discrimination between the two groups. The following features were selected: the average, total and variation of the nuclear fluorescence intensity per cell, the nuclear area, the skewness of the brightfield intensity distribution, the kurtosis of the brightfield intensity distribution and the total intensity of the brightfield within the regions of interest (ObjectAvgIntenCh1, ObjectTotalIntenCh1, ObjectVarIntenCh1, ObjectAreaCh1, ROI_SkewIntenCh3, ROI_KurtIntenCh3 and ROI_TotalIntenCh3 respectively). These parameters were used to generate 2 clusters using K-means clustering algorithm and then the percentage of injured cells per well was calculated, as previously described(21). We then compared the percentage of injured cells obtained to the total cell count in each well in order to detect cell lysis (ratio = % injured cells / cell count).

### 3. Large scale co-culture of clinical samples

We applied this strategy for the detection of SARS-CoV-2 in randomly chosen 104 anonymized nasopharyngeal swab samples. Initial RT-PCR ranged from 12 Ct to 34 Ct in 72 samples, and 32 samples with negative initial PCR were used as negative controls. Sample preparation and co-culture were performed as previously described(6). Briefly, 500 µl of the sample were processed into 0.22 µm pore sized centrifugal filter (Merck millipore, Darmstadt, Germany) and centrifuged at 12 000 g for 5 minutes. 50 µl were then inoculated on a monolayer of Vero E6 cells cultured in 96-well microplates. A negative control consisting of uninfected cells and a positive control consisting of cells infected with a 10^−4^ dilution of the IHUMI-3 strain were considered. A centrifugation step (700 x g for 1 h) was performed to enhance the entrance of the virus inside the cells. Plates were then incubated at 37°C and monitored for 7 days to search for cytopathic effects. In parallel, the same samples were processed using the classical isolation strategy based on the manual observation of cytopathic effects under an inverted microscope, in order to validate our strategy(6),(19),(22). For this strategy, co-cultures showing no cythopatic effects after 1 week were sub-cultured at day 7 and day 14 onto a fresh monolayer of cells for a complete observation of 3 weeks.

### 4. Results validation by scanning electron microscopy and RT-PCR

Positive co-cultures were processed with both scanning electron microscopy (SEM) and RT-PCR directly from culture supernatant to validate the presence of COVID-19 viral particles. Briefly, the SEM was performed using the SU5000 microscope (Hitachi High-Tech Corporation, Tokyo, Japan) allowing a rapid observation in about10 minutes without time consuming sample preparations(22). RT-PCR protocol was performed as previously described by Amrane *et al.* targeting the E gene(23). This RT-PCR was applied to wells showing a cytopathic effect to confirm that this effect was due to SARS-CoV-2 and to negative wells to confirm that the lack of cytopathic effect was not due to microscopically undetectable minimal viral growth.

### 5. Statistical analysis

The R Studio® and XLSTAT software were used to perform all statistical tests included in this paper.

### 6. Ethical statement

According to the procedures of the French Commission for Data Protection (Commission Nationale de l’Informatique et des Libertés), collected data were anonymized. The study was approved by the local ethics committee of IHU (Institut Hospitalo-Universitaire) - Méditerranée Infection (No. 2020-01).

## Results

### 1. Cytopathic effects and cell lysis detection

Figure 1 represents the fluorescence and brightfield images acquired with the cx7 microscope at days 1 and 6 post infection showing the early stages of infection of SARS-CoV-2 (Figure 1 - a, b) compared to advanced stages of infection and cell lysis (Figure 1 - g, h, i, j, k). Typical cytopathic effects consist of an increasing nuclear fluorescence intensity of the NucBlue fluorescent probe, in addition to nuclear fragmentation. These observations resulted in an increase in the average, total and variation intensity of the nucleus and a decrease in the nuclear area on the fluorescence images. Adding to this, infected cells become round and form aggregates resulting in an increasing total intensity, skewness and kurtosis on the brightfield images. Finally, advanced stages of infection are represented by cell lysis.

**Figure 1:**
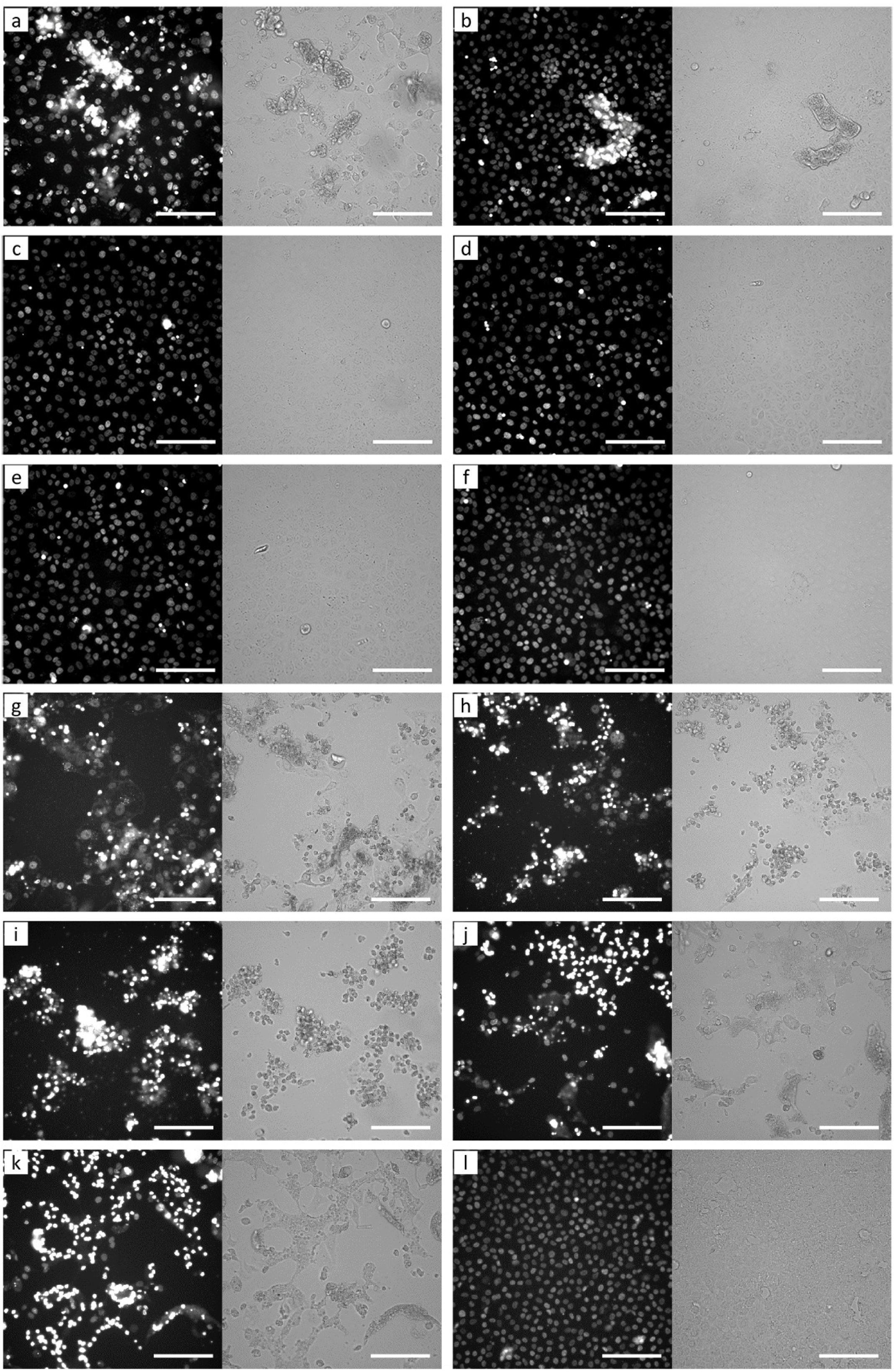
Kinetic monitoring of SARS-CoV-2 infection on Vero E6 cells over 6 days on the CX7 microscope showing cytopathic effects at different stages of infection. Images show respective fluorescence and brightfield images at different viral concentrations at days 1 and 6 post infection. Day 1: (a) Stock concentration, (b) 10^−1^ dilution, (c) 10^−2^ dilution, (d) 10^−3^ dilution, (e) 10^−4^ dilution and (f) negative control. Day 6: (g) Stock concentration, (h) 10^−1^ dilution, (i) 10^−2^ dilution, (j) 10^−3^ dilution, (k) 10^−4^ dilution and (l) negative control. Scale bars indicate 50 µm.

### 2. Automated detection results

The data extracted from the images were analyzed in the dedicated application in R Studio. The database of negative and positive controls served as training data for the clustering algorithm and a baseline of 2 to 3 % injured cells was predicted in the negative training data compared to a value of 50 to 55 % injured cells in the positive training data. The percentage of injured cells in each condition was predicted and then divided by the total cell count per well. This ratio allowed us to distinguish positive wells, showing cytopathic effects or cell lysis, from the negative control wells consisting of uninfected cells (Figure 2-a). Cytopathic effects were detectable up until the dilution 10^−4^ after 6 days of culture for the strain IHUMI-3 used in this study, which corresponds to the viral titer obtained by TCID50.

**Figure 2:**
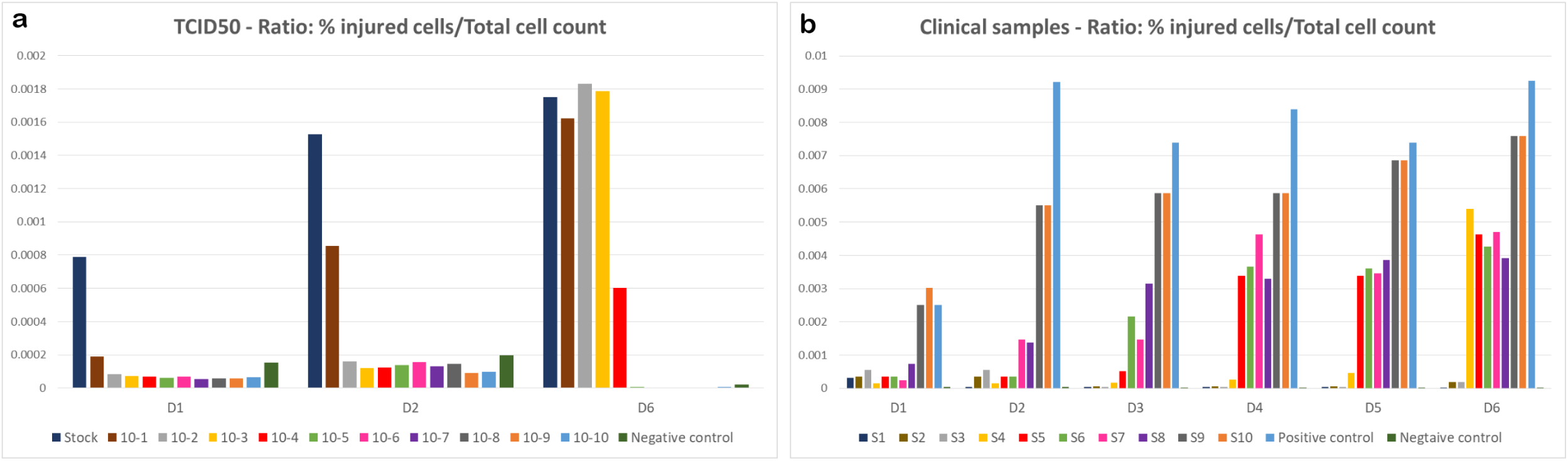
Automated detection of SARS-CoV-2 in co-culture. (A) Ratio of the percentage of injured cells on the total cell count of SARS-CoV-2 infected cells at different concentrations compared to the negative control over a period of 6 days. (B) Ratio of the percentage of injured cells on the total cell count of 10 clinical samples with different initial viral load over a period of 6 days. Initial viral load was negative in S1 and S2, 32 Ct in S3, 30 Ct in S4, 29 Ct in S5, 28 Ct in S6, 23 Ct in S7, 22 Ct in S8, 16 Ct in S9 and 15 Ct in S10.

Furthermore, the automation system allowed us to monitor co-culture on a daily basis without any intervention from the operators. The Momentum software was used to monitor the automation system linked to the HCS microscope. A screening process was predefined, thus allowing the proper incubation of the plates followed by the automated handling of the screening process at each specified time point.

### 3. Screening of clinical samples with the new HCS and the classic isolation strategies

Among the panel of 104 samples processed on the CX7 microscope, 32 samples had a negative initial PCR and were considered controls for system’s sensitivity, and therefore the corresponding co-cultures were negative. Among the remaining 72 samples, we managed to isolate the virus from 31 samples using our automated detection system. The detection delay ranged from 24 hours to 3 days for most samples and was prolonged to 6 days for samples with low viral load. Figure 2-b shows examples of co-culture results obtained with the automated detection system compared to the negative (uninfected cells) and positive (cells infected with the viral strain IHUMI-3) controls.

Regarding the classic isolation strategy, 30 viral strains were isolated from the tested panel of samples and the 32 samples with negative initial PCR had negative culture results as well. The majority of strains were recovered after fourth days of co-culture and only few were isolated at earlier stages. Three strains out of 30 were recovered after subcultures, 2 in the second week and 1 in the third week of co-culture.

A significantly higher percentage of positive samples was observed on a daily basis with the HCS strategy (Figure 3). Moreover, the majority of positive samples were isolated by the third day of co-culture using the HCS strategy, where 80 % positivity was obtained compared to only 26 % with the classic strategy (p value < 0.001).

**Figure 3:**
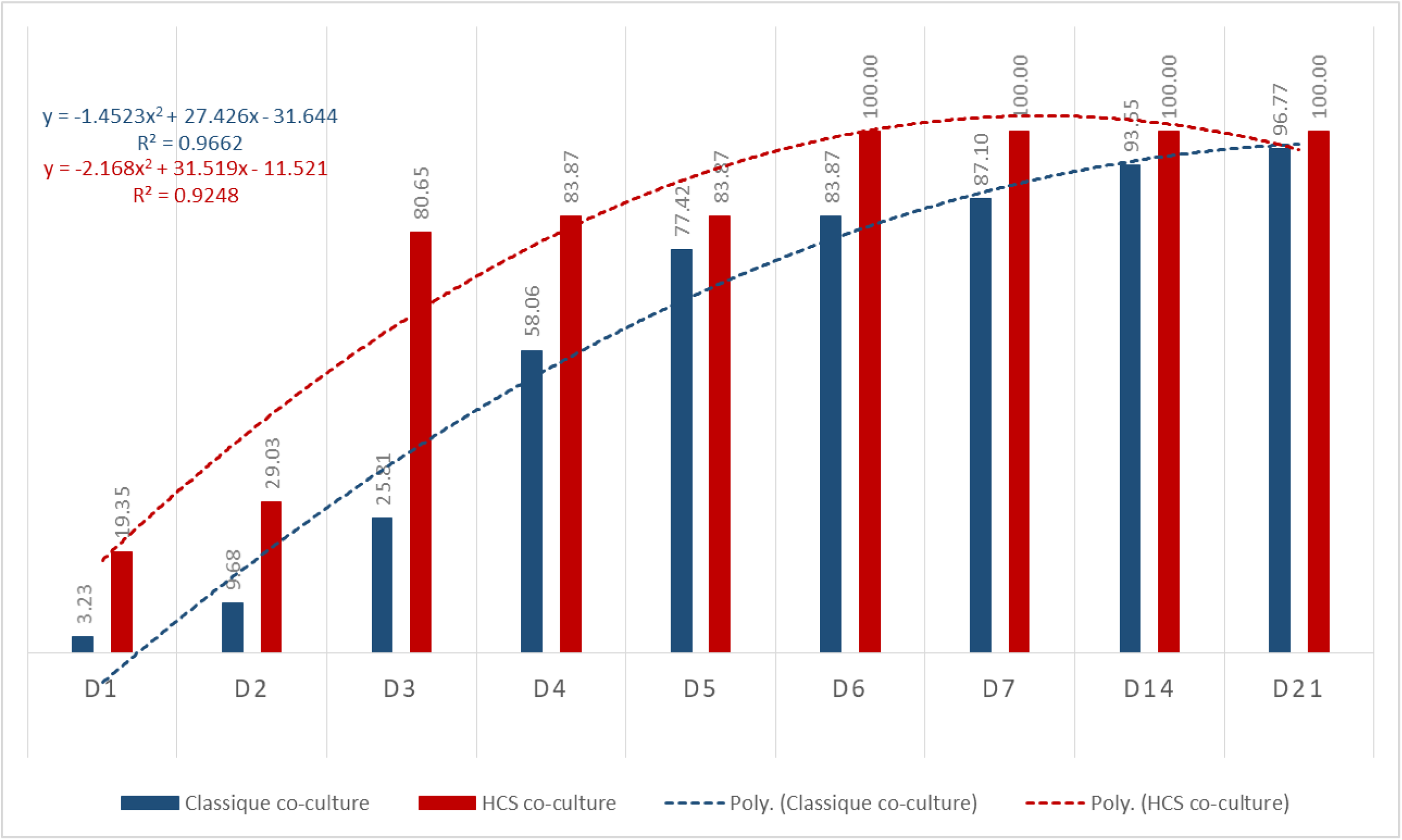
Cumulative percentage of isolated strains per day using the classic and the new HCS isolation strategies for samples detected as positive in co-culture. The dashed curve indicates the polynomial regression curve.

To validate our results, positive co-cultures were processed to scanning electron microscopy to confirm the presence of viral particles. We detected viral particles in the supernatant of all samples that were detected as positive by the HCS strategy. Figure 4 shows an example of particle detection in culture supernatant by SEM. RT-PCR performed on all wells correlated with the results of the microscopy-based detection. We then correlated the isolation rates obtained with both strategies to the initial viral RNA load (Figure 5). We obtained comparable isolation rates with the HCS isolation strategy, as compared with the classic strategy.

**Figure 4:**
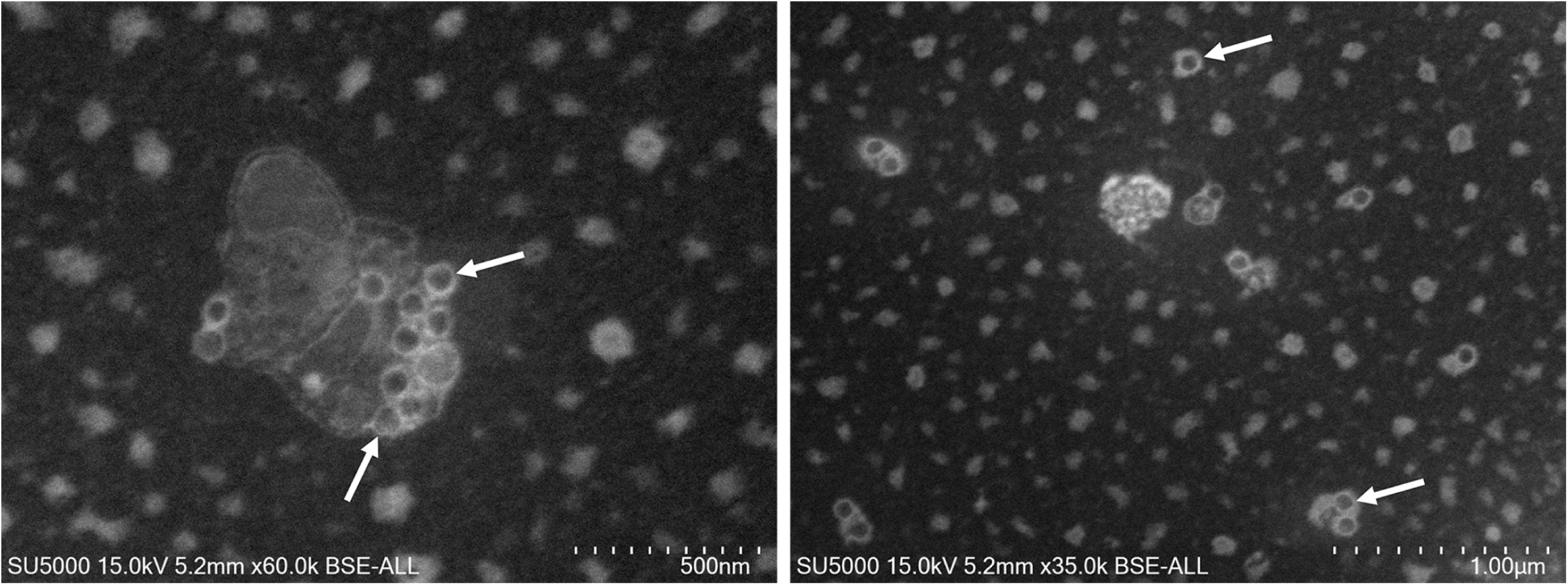
SEM images obtained with the SU5000 microscope showing SARS-CoV-2 particles isolated from clinical samples (white arrows). Acquisition settings and scale bars are generated on the original micrographs.

**Figure 5:**
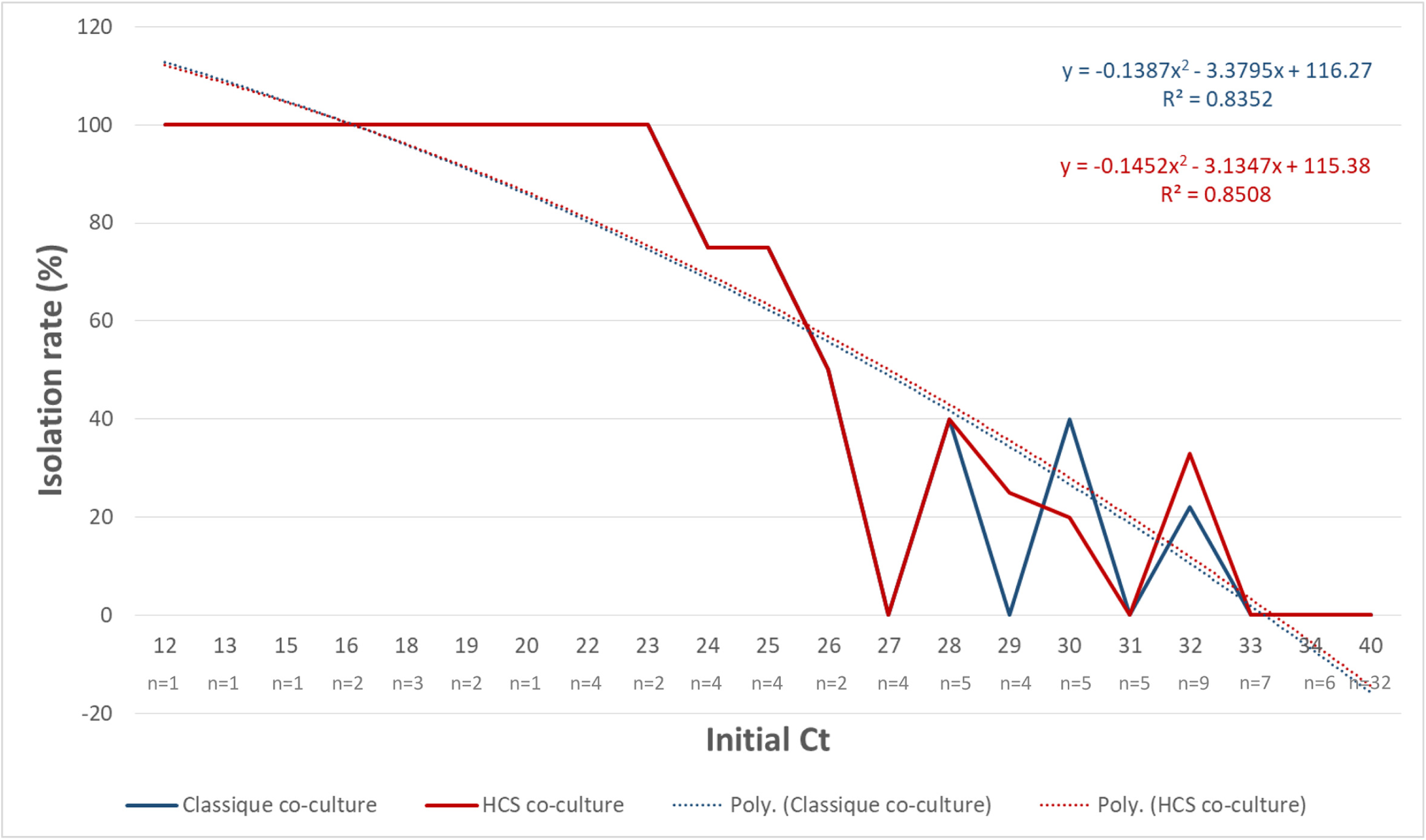
Isolation rate of SARS-CoV-2 from nasopharyngeal samples according to initial Ct values in samples (plain line) using the classic and the new HCS isolation strategies (40 Ct represents the samples with a negative initial PCR). The dashed curve indicates the polynomial regression curve.

Then, we correlated the isolation rates obtained with both strategies to the initial viral load (RT-PCR results) in each sample and the results are shown in figure 5. We obtained similar isolation rates with the HCS isolation strategy as well as with the classic strategy. Moreover, we noticed that most of the strains were recovered from samples with an initial viral load lower than 30 Ct with both strategies. Therefore, we calculated the positivity threshold of the isolation rate compared to the initial viral load in the samples using a ROC curve and we obtained a similar positivity threshold of 27 Ct for both isolation strategies.

## Discussion

Scientists are facing major challenges in the fight against Covid-19(15). Isolating the virus is a crucial factor, especially during pandemics, since all *in-vitro* analysis require having the virus(4). Furthermore, the greater the number of strains isolated, the better the understanding of the genetic diversity of this virus, especially since genome sequencing directly from samples is limited to viral load and a very poor genome assembly is obtained when the viral load is greater than 19 Ct (unpublished data). Developing an automated viral isolation technique was thus necessary to overcome the subjective and time consuming manual microscopic observations. In this work, we were able to co-culture a large amount of clinical samples and monitor them with a fully automated system, which reduced the workload and time required from laboratory technicians. The main advantage of this technique is the automation as it allows limiting the risk of exposure or contamination of the personnel, since plate monitoring and data analysis could be carried out from distance, thus avoiding direct contact and manual observations of co-cultures. Similar isolation rates were obtained with both isolation strategies, which validated the efficiency of our new automated system. Moreover, this isolation rate was obtained in 1 week with the HCS strategy without any subcultures, contrary to the classic technique with weekly subcultures for a complete incubation time of 3 weeks. Subsequently, since the loss of cultivability of the virus in samples allows to consider patients at low risk of contamination, it helps in the decision to discharge them from infectious diseases wards(19). Using this HCS isolation strategy allows us to answer this question in one week. This is especially critical at the beginning of an epidemic or when PCR detection systems have to be modified. Thus, this new automated isolation strategy is applicable during the current crisis to recover strains from suspected samples in a safe and rapid way. Further work is underway to use this technique for large-scale drug susceptibility testing of SARS-CoV-2 strains isolated from patients. And, finally, the algorithms used here could be adapted and applied for the detection and isolation of other viruses from clinical samples in case of known and emerging viral diseases.

## Acknowledgments

We sincerely thank Takashi Irie, Kyoko Imai, Shigeki Matsubara, Taku Sakazume, Toshihide Agemura, Yusuke Ominami, Hisada Akiko and the Hitachi team in Japan Hitachi High-Tech Corporation, Toranomon Hills Business Tower, 1-17-1 Toranomon, Minato-ku, Tokyo 105-6409, Japan) for the collaborative study conducted together with IHU Méditerranée Infection, and for the installation of a SU5000 microscope at the IHU Méditerranée Infection facility.

